# pOPARA: Vectors for Golden Gate assembly of expression constructs containing the araBAD promoter

**DOI:** 10.1101/2023.05.31.543011

**Authors:** Lesley Milnes, Mark Youles, Josephine H. R. Maidment

## Abstract

Heterologous protein production is often required to investigate the structural, biochemical, and biophysical properties of a protein of interest. Frequently, optimisation of expression conditions is required to obtain soluble protein and maximise yield. Trialling a variety of solubility and purification tags, as well as constructs containing different regulatory elements, is desirable. Golden Gate cloning allows modular assembly of different constructs using Type IIS restriction enzymes. The pOPIN vector suite, which utilises the T7 expression system, has been adapted to be compatible with Golden Gate assembly. Here, we present the pOPARA vectors. Expression from pOPARA vectors is driven by the araBAD promoter (pBAD) and is induced by addition of arabinose to the culture medium. pOPARA allows modular assembly of expression constructs using Golden Gate cloning with the CDS of interest and an optional C-terminal tag. pOPARA1 contains a carbenicillin resistance cassette flanked by restriction sites to allow exchange of the selectable markers. In pOPARA2, the carbenicillin resistance cassette has been exchanged for a spectinomycin resistance cassette. We demonstrate that both vectors can be used to express and produce a control protein.

## Introduction

Production of recombinant proteins in heterologous expression systems is central to many studies seeking to characterise protein activity, function, interactions, and structure. The bacterium *E. coli* is a well-established heterologous expression system and frequently the first system used by researchers attempting to produce a new protein of interest (Rosano and Ceccarelli, 2014). However, obtaining enough soluble protein for downstream analyses can require extensive screening of expression constructs and bacterial growth conditions.

Bacterial expression constructs typically contain the coding sequence of the protein of interest under the control of an inducible promoter to enable timely induction of protein production, generally during the log phase of bacterial growth. The *lac* promoter, from the well-studied *lac* operon (Jacob and Monod, 1961), has been extensively used in bacterial expression vectors. Gene expression from this promoter is induced by addition of the allolactose mimic isopropyl β-d-1-thiogalactopyranoside (IPTG) to the culture medium. In the absence of IPTG, the *lac* repressor LacI blocks binding of RNA polymerase to the *lac* promoter. IPTG binds to LacI and the resulting allosteric changes lead to the dissociation of LacI from the DNA, thus allowing RNA polymerase to initiate transcription. The pUC vectors (Vieira and Messing, 1982) contain the *lac* promoter, while the pTrc vectors (Amann et al., 1988) contain a *tac* promoter (de Boer et al., 1983), a synthetic hybrid of the *lac* and *trp* promoters which drives higher expression levels than the *lac* promoter. In the pET (Dubendorff and Studier, 1991) and pOPIN (Berrow et al., 2007) vectors, the CDS is cloned downstream of the phage T7 promoter which is recognised by the phage T7 RNA polymerase (Dubendorff and Studier, 1991, Studier et al., 1990, Rosenberg et al., 1987). The coding sequence for this polymerase, in turn, is placed under the control of the lac promoter, such that addition of IPTG to the culture medium induces expression of the T7 RNA polymerase, which subsequently drives high expression of the gene of interest.

The *araBAD* promoter of the *E. coli araBAD* operon has also been used in bacterial expression vectors (Guzman et al., 1995, Marschall et al., 2017). The *araBAD* operon contains three structural genes (*araB, araA* and *araD*), transcribed as a single transcript, encoding enzymes which collectively catalyse the conversion of L-arabinose to D-xylulose-5-phosphate (Schleif, 2000). The *araBAD* promoter is regulated by L-arabinose concentration through allosteric effects of L-arabinose on the regulator AraC (Sheppard and Englesberg, 1967, Englesberg et al., 1965). The gene encoding AraC is located upstream of the *araBAD* operon and is transcribed from a separate promoter. In the absence of L-arabinose, an AraC homodimer binds to two DNA sites upstream of the *araBAD* operon, leading to formation of a DNA loop which blocks binding of RNA polymerase to the *araBAD* promoter, inhibiting expression of the structural genes. When L-arabinose is present, the AraC homodimer undergoes a conformational change and switches DNA binding sites, removing the DNA loop and allowing RNA polymerase to bind to the *araBAD* promoter and initiate transcription (Figure 1). The pBAD series of expression vectors (Guzman et al., 1995) contain the *araBAD* operon, where the *araBAD* structural genes have been replaced with a multiple cloning site (MCS).

**Figure 1.**
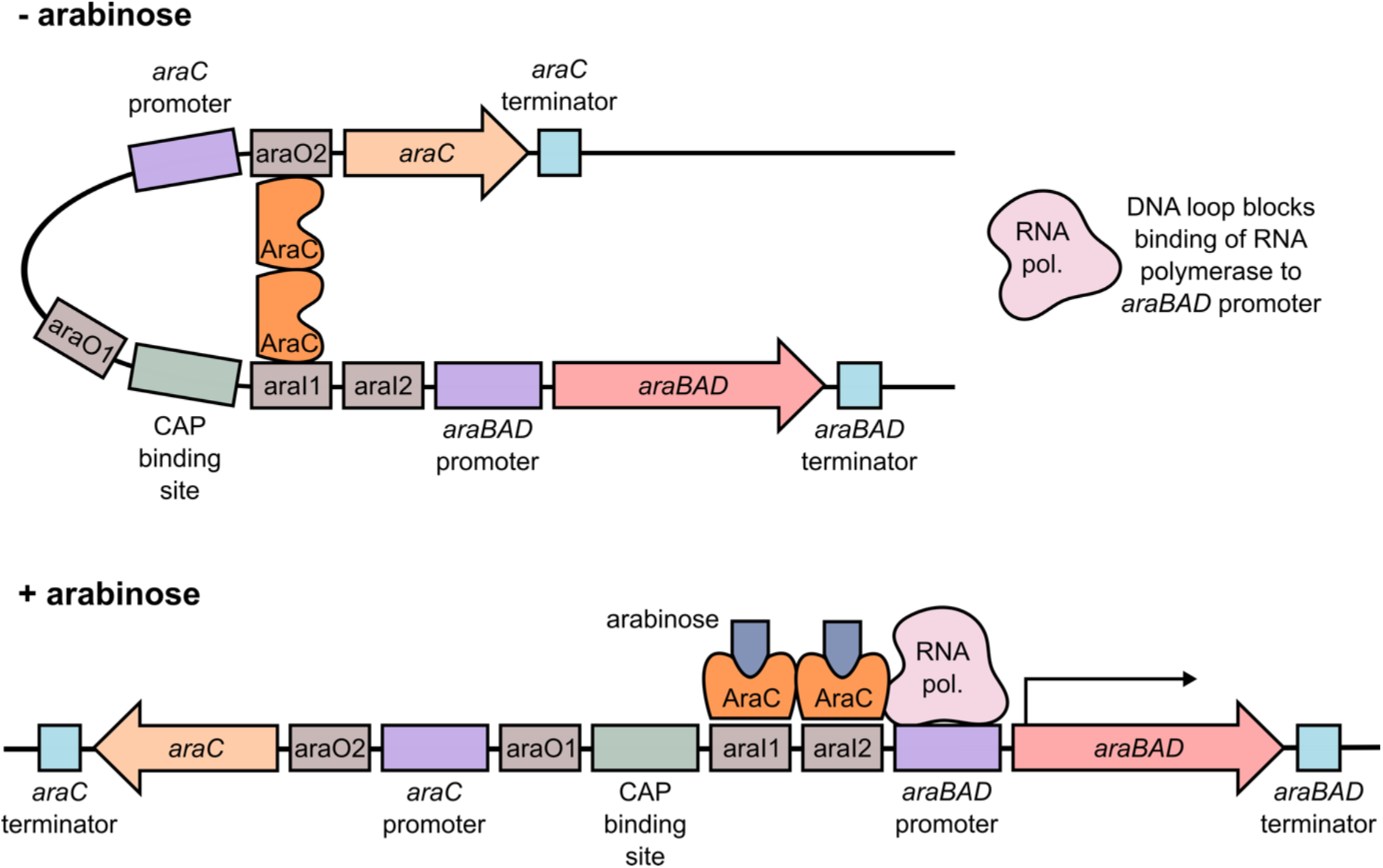
Diagram of the bacterial *araBAD* operon. In the absence of L-arabinose, the dimeric repressor protein AraC binds to two sites, forming a DNA loop which blocks the binding of RNA polymerase to the *araBAD* promoter. In the presence of L-arabinose, AraC undergoes conformational changes and switches DNA binding sites. This removes the DNA loop, enabling RNA polymerase to bind to the *araBAD* promoter and initiate transcription of the *araBAD* genes.

Different promoters vary in both the basal level of expression (in the absence of an inducing molecule) and the maximal level of expression achieved following induction. The *lac* promoter is notoriously “leaky”, with detectable basal expression of the gene of interest in the absence of IPTG (Nielsen et al., 2007). This can be attributed to insufficient LacI repressor to occupy all available binding sites, and is exacerbated by the use of high copy number plasmids (Rosano and Ceccarelli, 2014). This “leaky” expression can be reduced by using a mutated version of the *lacI* promoter which supports higher expression of *lacI* (Penumetcha et al., 2010). For plasmids using the T7 system, pLysS or pLysE plasmids produce T7 lysozyme, an inhibitor of T7 RNA polymerase, and use of bacterial hosts containing these plasmids can counter “leaky” expression of T7 RNA polymerase (Studier, 1991). However, for proteins which are toxic to *E. coli*, even low levels of background expression can be problematic. When compared to the *lac* promoter, the *araBAD* promoter results in significantly lower basal expression in the absence of L-arabinose (Nielsen et al., 2007). Further repression of gene expression from the *araBAD* promoter can also be achieved by addition of glucose to the bacterial culture (Lee and Jung, 2007).

Expression from *lac* or T7 promoters is often favoured due to high expression levels of the gene of interest upon induction. However, such high expression can be problematic when the protein of interest has a deleterious effect on bacterial growth or is toxic to *E. coli*. Very high expression can increase protein misfolding and accumulation in inclusion bodies. The *araBAD* promoter typically supports lower levels of expression than the *lac* or T7 promoters, and expression can be modulated by varying the concentration of L-arabinose used for induction and/or adding glucose to the culture following induction (Lee and Jung, 2007, Siegele and Hu, 1997). Therefore, for certain proteins of interest, use of the *araBAD* promoter may improve soluble protein yield. As the properties of different promoters affects their suitability for production of a particular protein of interest, it is useful to screen expression constructs containing different promoters.

Golden Gate cloning exploits the property of Type IIS restriction endonucleases to cut unspecific DNA sequence at a specific location outside their recognition site to assemble multiple DNA fragments in a single digestion-ligation (dig-lig) reaction (Engler and Marillonnet, 2014, Weber et al., 2011, Engler et al., 2008). Golden Gate cloning has several advantages over restriction cloning. DNA cleavage by the Type IIS restriction endonucleases *Bsa*I and *Bpi*I results in four nucleotide overhangs, the sequence of which can be selected by the user to allow either scarless assembly or assembly with only a very short scar. This is particularly advantageous when assembling constructs for protein production, where additional amino acids can significantly impact protein function. A single enzyme can release DNA fragments with distinct overhangs, which can then be assembled such that the restriction sites are absent in the ligated product. This prevents digestion of the ligated product, allowing both digestion and ligation to take place in a single reaction, reducing the number of steps in the cloning pipeline. Finally, Golden Gate cloning supports modular assembly of expression constructs (Weber et al., 2011). Plasmids containing coding sequences, promoters, terminators and purification or solubility tags (referred to as “level 0” parts) can be assembled into “level 1” expression cassettes, which can subsequently be combined into “level 2” multigene constructs.

The pOPIN-GG vectors (Bentham et al., 2021) represent a highly useful resource for modular assembly of expression constructs. These vectors were adapted from the pOPIN vector suite, generated by the Oxford Protein Production Facility (OPPF, now PPUK; (Berrow et al., 2007)), which uses the T7 promoter system to drive expression.

Here, we present the expression vectors pOPARA1 and pOPARA2, which have been adapted from a pBAD vector (Guzman et al., 1995) to enable modular Golden Gate assembly of expression constructs where expression is driven by the *araBAD* promoter. Additionally, pOPARA1 contains a carbenicillin resistance cassette which can be exchanged for other resistance cassettes in a digestion-ligation reaction with the Type IIS restriction enzyme *Bsm*BI. We show that both pOPARA1 and pOPARA2 drive arabinose-inducible expression of a gene for soluble protein production.

## Results

### Design of pOPARA1

We used the pBAD24 vector sequence (Guzman et al., 1995) as the basis for pOPARA1. pBAD24 contains a multiple cloning site (MCS) between the *araBAD* promoter and the *rrnB* terminator. We replaced the MCS for a red fluorescent protein (RFP) cassette, as a negative selection marker, flanked by *Bsa*I restriction sites, positioned such that digestion of the vector with *Bsa*I would produce 5’ GCTT and 3’ AATG overhangs. The overhangs were selected to follow the standard syntax for Type IIS restriction endonuclease-mediated assembly defined and adopted by the plant synthetic biology community (Patron et al., 2015). pOPARA1 can therefore be used to produce either C-terminally tagged or untagged proteins. For untagged proteins, *Bsa*I digestion of a plasmid or PCR product containing the CDS should generate 5’ AATG and 3’ GCTT overhangs, where the ATG of the 5’ AATG overhang is the start codon and the 3’ GCTT overhang follows a stop codon. For C-terminally tagged proteins, *Bsa*I digestion should produce 5’ AATG and 3’ TTCG overhangs, where again the ATG of 5’ AATG is the start codon, and the TCG of 3’ TTCG is a codon for serine (Patron et al., 2015). *Bsa*I digestion of the plasmid or PCR product encoding the C-terminal tag should generate 5’ TTCG and 3’ GCTT overhangs, where the TCG of 5’ TTCG is a codon for serine, and the 3’ GCTT overhang follows a stop codon. Multiple level 0 modules containing C-terminal tags are already available on Addgene. By following the standard syntax outlined in Patron et al. (Patron et al., 2015), the same CDS and C-terminal tag modules can be used for assembly of constructs in pOPIN-GG and pOPARA vectors (Figure 2), facilitating comparison of the different expression induction systems for a protein of interest.

**Figure 2.**
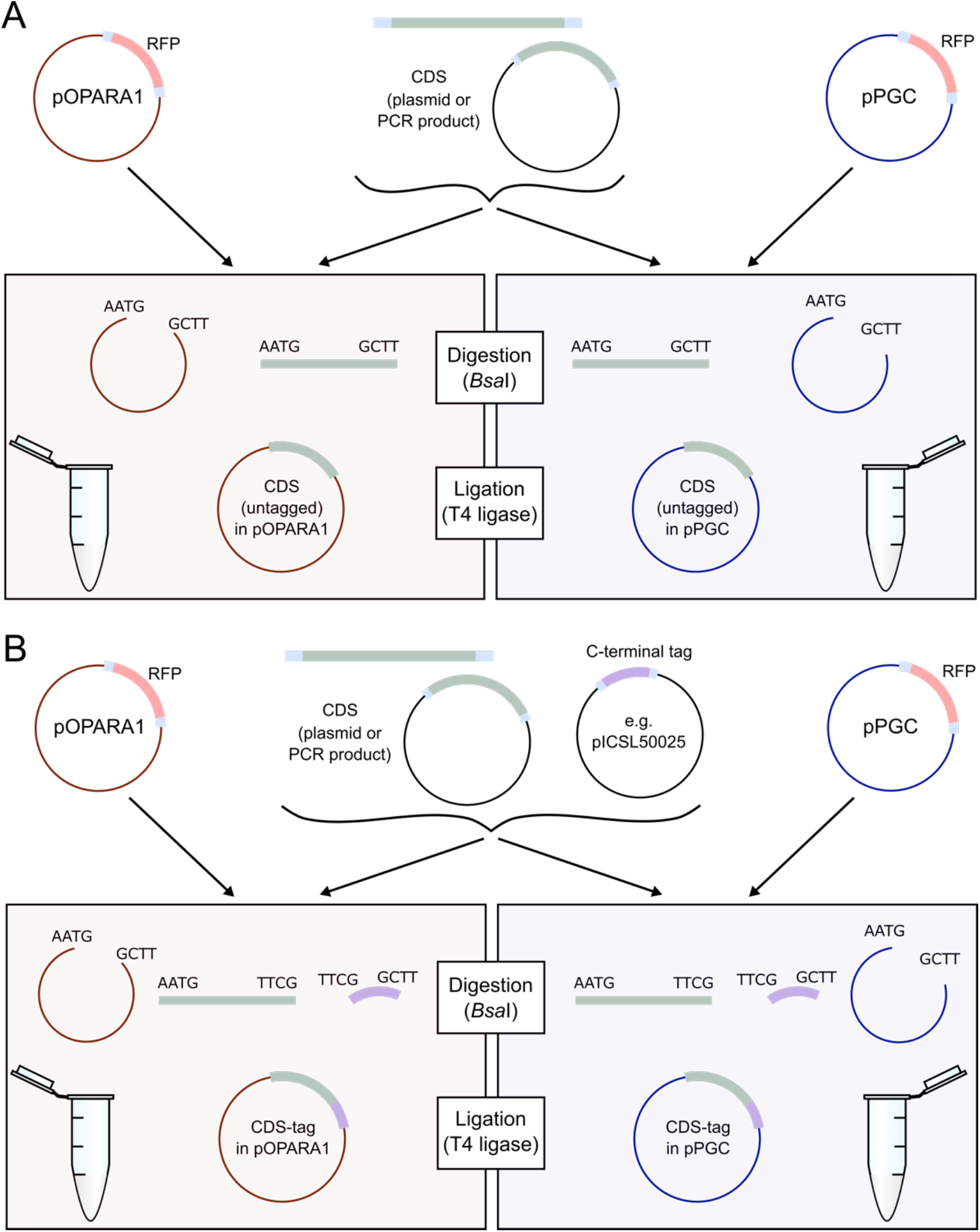
Diagram of the workflow to assemble pOPARA1 expression constructs by Golden Gate cloning. Digestion with *Bsa*I of pOPARA1 or pPGC produces 3’ AATG and 5’ GCTT overhangs. The same CDS and C-terminal tag modules can be assembled into pOPARA and pOPIN-GG (pPGC) vectors to produce untagged or C-terminally tagged proteins. **A**. To generate an expression construct for an untagged protein, digestion with *Bsa*I of either a level 0 module or PCR product containing the CDS should produce the overhangs 5’ AATG and 3’ GCTT. **B**. For C-terminal tagging, digestion with *Bsa*I of either a level 0 module or PCR product containing the CDS should produce the overhangs 5’ AATG and 3’ TTCG, and digestion with *Bsa*I of a level 0 module containing the C-terminal tag sequence should produce overhangs 5’ TTCG and 3’ GCTT.

Production and purification of protein complexes from *E. coli* can be achieved by co-transformation of E. coli with two plasmids, each containing a gene for one of the proteins and each containing a different antibiotic resistance gene to allow selection of cells containing both plasmids. Like pBAD24, pOPARA1 contains the *bla* gene (encoding beta-lactamase) which confers resistance to carbenicillin. To facilitate production of pOPARA vectors with different antibiotic resistance genes, we introduced *Bsm*BI restriction sites on either side of the carbenicillin resistance cassette. Digestion of pOPARA1 with *Bsm*BI releases the carbenicillin resistance gene (including the promoter and terminator) and the resulting backbone has overhangs 5’ CGCG and 3’ TACA.

The native pBAD24 plasmid sequence contains two *Bsa*I restriction sites and one *Bsm*BI site. A single nucleotide substitution was introduced into each site to avoid unwanted cuts during digestion-ligation reactions (see Materials & Methods). The final pOPARA1 plasmid was commercially synthesised by Gene Universal Inc. (Newark, DE). A schematic representation of pOPARA1 is shown in Figure 3. The annotated sequence of pOPARA1 is provided in Supplementary Data 1 and the plasmid is available from Addgene (ID #200416).

**Figure 3.**
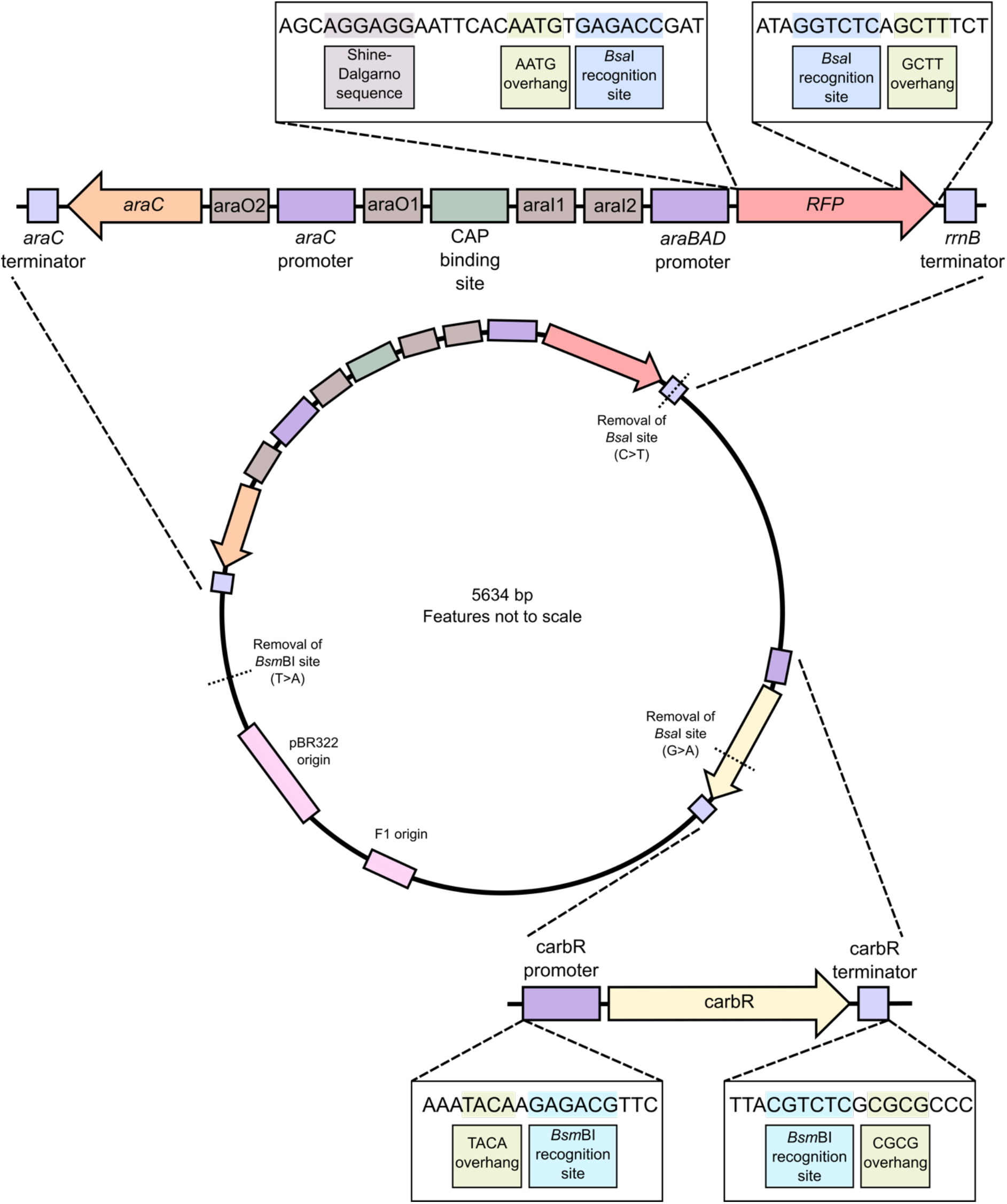
Schematic representation of pOPARA1. pOPARA1 contains the *araBAD* operon, where the *araBAD* structural genes have been replaced by a red fluorescent protein (RFP) cassette flanked by *Bsa*I restriction sites. The carbenicillin resistance cassette is flanked by *Bsm*BI restriction sites. Highlighted plasmid features are not to scale.

### BsmBI-mediated exchange of the selection cassette of pOPARA1 to generate pOPARA2

pAGM1299 (Addgene plasmid # 47988; (Weber et al., 2011)) contains the *aadA* gene (encoding an adenylyltransferase) which mediates resistance to spectinomycin (and streptomycin). The spectinomycin resistance gene, including the promoter and terminator regions, was PCR-amplified from pAGM1299 to give a PCR product flanked by *Bsm*BI restriction sites. Digestion of the PCR product with *Bsm*BI results in 5’ TACA and 3’ CGCG overhangs. The primers used to amplify the spectinomycin resistance gene from pAGM1299 are in Table 1. pOPARA2 was generated via an *Bsm*BI digestion-ligation reaction between pOPARA1 and the pAGM1299 PCR product (see Materials and Methods). A similar approach can be used to exchange the carbenicillin resistance gene of pOPARA1 for other antibiotic resistance genes (e.g. *aphA1* for kanamycin resistance, *catA1* for chloramphenicol resistance) as required by the user. It should be noted that the *Bsm*BI restriction sites are removed in the digestion-ligation reaction, so further exchange of selection cassettes is not possible. The workflow and a schematic representation of pOPARA2 are shown in Figure 4. The annotated sequence of pOPARA2 is provided in Supplementary Data 2 and the plasmid is available from Addgene (ID #200417).

**Table 1.**
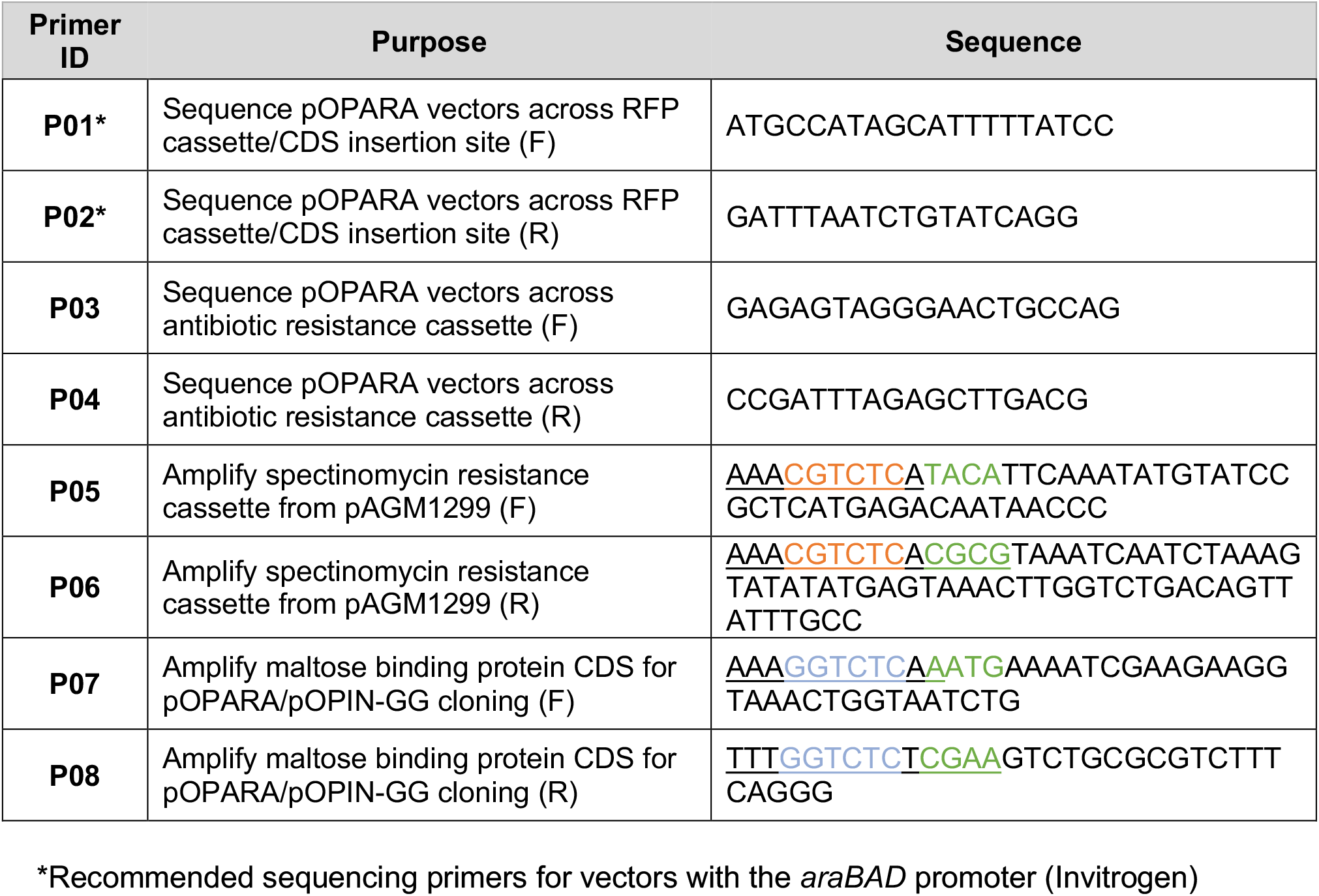
Primers used for PCR amplification and sequencing. For primers P05-P12, the flanking sequence is underlined. Introduced *Bsm*BI and *Bsa*I sites are indicated in orange and blue, respectively. Overhangs revealed by restriction digestion with *Bsm*BI or *Bsa*I are indicated in green.

**Figure 4.**
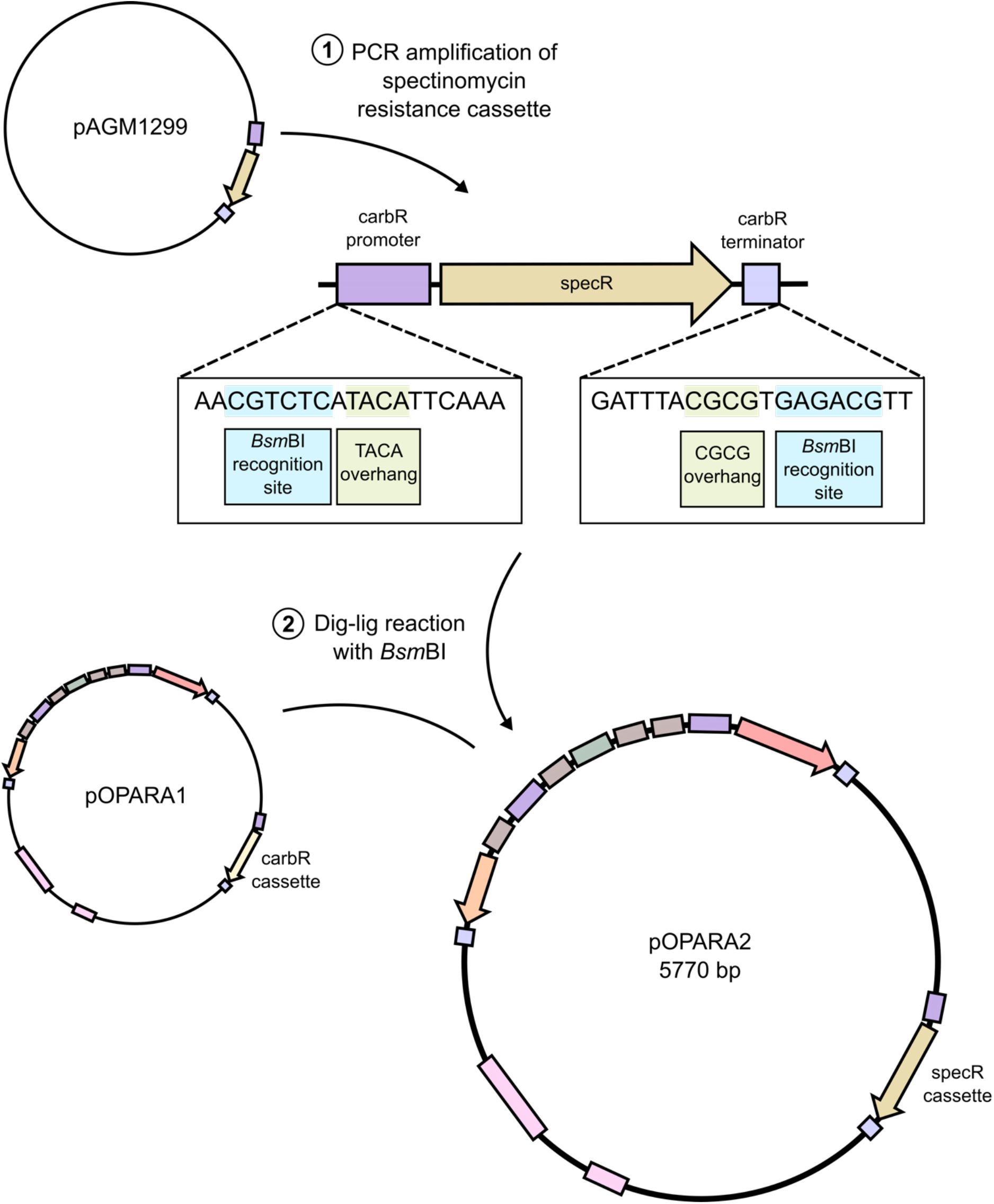
Diagram of the workflow used to generate pOPARA2. The spectinomycin resistance cassette was PCR-amplified from pAGM1299 and exchanged for the carbenicillin resistance cassette of pOPARA1 via a digestion-ligation reaction using the Type IIS restriction enzyme *Bsm*BI.

### pOPARA1 and pOPARA2 enable protein production

To test whether pOPARA1 and pOPARA2 support recombinant protein production in an arabinose-dependent manner, we cloned *E. coli* maltose binding protein (MBP, (Duplay et al., 1984)) with a C-terminal 6xHis tag, into pOPARA1, pOPARA2 and the pOPIN-GG vector pPGC (Bentham et al., 2021). The resulting plasmids were used to transform *E. coli* BL21(DE3) cells.

Transformed cells were grown in L media and expression was induced using either 0.2% L-arabinose (pOPARA1 and pOPARA2) or 1mM IPTG (pOPIN-GG). Proteins were purified from the soluble fraction of the cell lysate by Ni^2+^-immobilised metal affinity chromatography (IMAC). Analysis of the eluate by SDS-PAGE showed that MBP was produced from both pOPARA1 and pOPARA2 (Figure 5). We observed reduced protein yield from pOPARA1 and pOPARA2 compared to pOPIN-GG. We demonstrate here that the pOPARA vectors support production of a control protein.

**Figure 5.**
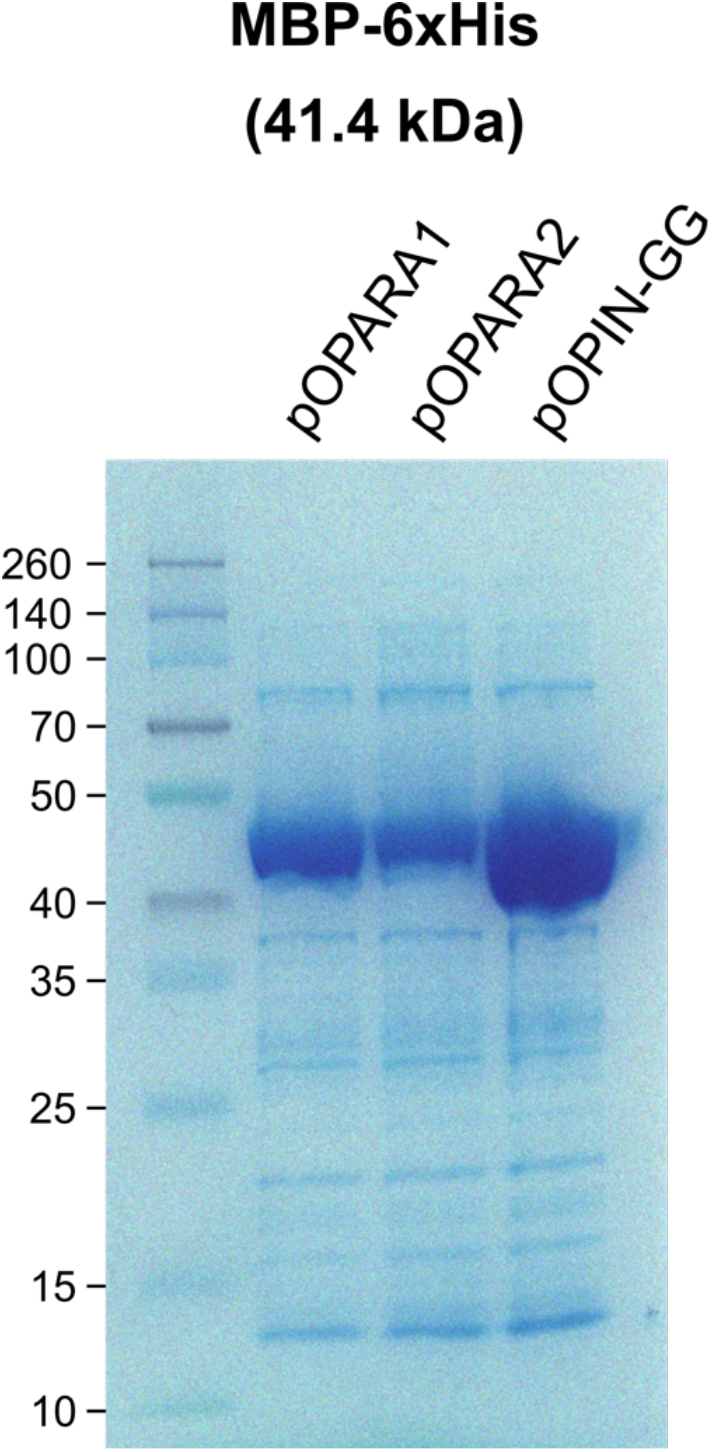
Production of a control protein using pOPARA1 and pOPARA2. SDS-PAGE gel of *E. coli* maltose binding protein (MBP) with a C-terminal 6xHis tag produced from pOPARA1, pOPARA2 or the pOPIN-GG vector pPGC and purified by immobilised metal affinity chromatography.

## Discussion

Production and purification of recombinant proteins is a requirement for many studies in the field of protein biochemistry. Obtaining sufficient soluble protein is often a bottleneck for such studies and requires optimisation of protein production conditions. Screening of different expression constructs, containing different solubility and/or purification tags and different promoters, is desirable. The ability to rapidly, and in parallel, assemble multiple constructs facilitates high-throughput screening and the identification of optimal conditions for production of a protein of interest. Golden Gate cloning, which exploits the property of Type IIS restriction enzymes to cut outside their recognition sequence, allowing digestion and ligation reactions to proceed simultaneously without the risk of further digestion of the ligated product, is a popular method for high-throughput construct assembly. Golden Gate “toolboxes” have already been developed to assemble constructs for yeast (Prielhofer et al., 2017), baculovirus (Spielvogel et al., 2021, Neuhold et al., 2020), and plant (Engler et al., 2014) transformation. Previously, the pOPIN-GG vectors (Bentham et al., 2021) were modified from the pOPIN vector suite to support modular assembly of *E. coli* expression constructs using Golden Gate cloning. Expression from the pOPIN and pOPIN-GG vectors is driven by the T7 promoter. Here, we present the pOPARA vectors, which similarly allow Golden Gate assembly of expression constructs, but with expression under the control of the *araBAD* promoter.

As the choice of promoter can affect the yield of soluble protein, testing different promoters is desirable when screening expression conditions. The *lac* promoter is generally “leakier” than *araBAD* promoter, with higher levels of basal expression in the absence of an inducing molecule (Nielsen et al., 2007). For proteins which have deleterious effect on bacterial growth or survival, using the *araBAD* promoter may increase protein yield. Following induction, the *araBAD* promoter typically supports lower levels of expression than the lac or T7 promoters. We also observed reduced protein yield from pOPARA1 and pOPARA2 compared to pOPIN-GG. However, in some cases, very high levels of protein production can lead to protein misfolding and accumulation of the protein in inclusion bodies. Furthermore, expression from the araBAD promoter can be controlled by varying the concentration of L-arabinose used for induction (Siegele and Hu, 1997), which can be valuable when tight control of expression levels is required.

Like the pOPIN-GG vectors, the pOPARA vectors follow the standard syntax for Golden Gate parts agreed by the synthetic biology community (Patron et al., 2015). Crucially, this means that the same CDS and C-terminal tag modules can be incorporated into both pOPIN-GG and pOPARA vectors in parallel Golden Gate reactions. The modularity of this system allows new purification and solubility tags to be introduced into both pOPIN-GG and pOPARA vectors as they become available.

We designed pOPARA1 such that digestion with *Bsa*I gives 3’ AATG and 5’ GCTT overhangs. In accordance with the syntax for Golden Gate parts (Patron et al., 2015), pOPARA1 can therefore either accept a single CDS (for untagged proteins) or a CDS and C-terminal tag. A pOPARA vector which could accept a N-terminal tag and a CDS would be a useful addition to the vector suite, as the position of a tag can affect the amount of soluble protein obtained. By exchanging the AATG overhang at the 5’ end of the RFP cassette for CCAT, a pOPARA vector could be produced for N-terminal tagging.

The carbenicillin resistance cassette of pOPARA1 is flanked by *Bsm*BI restriction sites, such that a digestion-ligation reaction with the type IIS restriction enzyme *Bsm*BI can facilitate exchange of the resistance cassette. We successfully demonstrated this by exchanging the carbenicillin resistance cassette of pOPARA1 for the spectinomycin resistance cassette of pAGM1299, generating pOPARA2. We propose that co-transformation of *E. coli* with pOPARA1 and pOPARA2 can be used for production and purification of protein complexes. We note that both pOPARA1 and pOPARA2 share the same origin of replication. While it is generally advised to co-transform *E. coli* with plasmids containing different origins of replication (from different compatibility groups, (Kerrigan et al., 2011)), to promote maintenance of both plasmids in the transformed cells, it has been reported that the selection pressure imposed by addition of the two antibiotics to the culture medium is sufficient to ensure maintenance of both plasmids in *E. coli* cultures for the time required for induction of gene expression and protein production (Yang et al., 2001). Consistent with this, there are multiple examples of protein complexes produced from co-transformation of *E. coli* with incompatible plasmids (Maidment et al., 2021, De la Concepcion et al., 2018, Maqbool et al., 2016).

In summary, the pOPARA vectors enable modular assembly of *E. coli* expression constructs where expression is driven by the tightly regulated *araBAD* promoter. The same CDS and C-terminal tag modules can be incorporated into both pOPIN-GG and pOPARA vectors, facilitating comparison of the two promoters for a protein of interest. We have made the pOPARA1 and pOPARA2 vectors available on Addgene (ID #200416 and #200417).

## Materials and Methods

### Design of pOPARA1

The MCS of pBAD24 (Guzman et al., 1995) was exchanged for the red fluorescent protein (RFP) cassette from pPGC-C (Addgene plasmid # 174580,(Bentham et al., 2021)) flanked by *Bsa*I restriction sites, positioned such that digestion of the vector with *Bsa*I would produce 5’ GCTT and 3’ AATG overhangs. pBAD24 contains Shine-Dalgarno (SD) and Kozak sequence motifs, which serve as ribosomal binding site in bacterial mRNA and the protein translation initiation site in eukaryotic mRNA, respectively. The SD sequence (AGGAGG) is located 8 nucleotides upstream of the ATG start codon, and this position relative to the start codon was maintained in pOPARA1. The Kozak sequence (ACCATG, where ATG is the start codon) could not be maintained due to the requirement to introduce the AATG overhang. As this vector is expected to be used for expression in bacterial, rather than eukaryotic, hosts, we decided to prioritise retaining the AATG overhang (in line with the standard syntax described in (Patron et al., 2015)) over the consensus Kozak sequence.

*Bsm*BI restriction sites were inserted immediately upstream of the promoter and immediately downstream of the terminator of the carbenicillin resistance cassette. The four nucleotides immediately upstream of the promoter (TACA) and the four nucleotides immediately downstream of the terminator (CGCG) prior to insertion of the *Bsm*BI sites were selected as the overhangs. As *Bsm*BI cuts one nucleotide downstream of the CGTCTC recognition site, an additional nucleotide was added such that digestion of the plasmid with *Bsm*BI would excise the carbenicillin resistance cassette and the backbone would have 5’ CGCG and 3’ TACA overhangs.

The region of the pBAD24 plasmid sequence used for pOPARA1 contains two *Bsa*I restriction sites and one *Bsm*BI site. To domesticate (remove restriction sites from) pOPARA1 to prevent unwanted restriction digestion, a single nucleotide substitution was made in each of the restriction sites. For the *Bsa*I site located in the CDS of the carbenicillin resistance gene *bla*, the G > A substitution is synonymous (does not affect the amino acid sequence of the protein product). Details of the domestication substitutions are in Table 2.

**Table 2.** Nucleotide substitutions introduced to domesticate pOPARA1.

| Restriction site | Nucleotide | Location | Substitution |
| --- | --- | --- | --- |
| *Bsal* | 2614 | *rrnB* terminator | C > T (GGTCTC > GGTTTC) |
| *Bsal* | 3686 | *bla* CDS | G > A (GGTCTC > GATCTC) |
| *BsmBI* | 5367 | - | T > A (CGTCTC > CGACTC) |

### *BsmBI*-mediated exchange of the selection cassette of pOPARA1 to generate pOPARA2

The spectinomycin resistance cassette (promoter, *aadA* CDS and terminator) was PCR-amplified from pAGM1299 (Addgene plasmid # 47988; (Weber et al., 2011)) using Phusion high-fidelity DNA polymerase (New England Biolabs Ltd) according to the manufacturer’s instructions. Primers P05 and P06 (Table 1) were designed to introduce flanking overhangs (5’ TACA and 3’ CGCG) and *Bsm*BI restriction sites into the PCR product.

The amplicon was combined with 100 ng pOPARA1 in a 2:1 molar ratio and the DNA mixture was brought to a total volume of 10 µl with dH_2_O. This was combined with 1.5 µl 10X T4 DNA ligase buffer (New England Biolabs Ltd.), 1.5 µl 10 mg/ml bovine serum albumin (BSA; New England Biolabs Ltd.), 1 µl T4 DNA ligase (2000U; New England Biolabs Ltd.) and 1µl *BsmB*I-v2 (10U; New England Biolabs Ltd). *BsmB*I-v2 is an isoschizomer of *Esp*3I. Thermocycling conditions for the digestion-ligation reaction were 30 cycles of (1 minute at 42 °C, 1 minute at 16 °C) followed by 5 minutes at 60 °C. 5 µl reaction product was used to transform chemically competent *E. coli* DH5α cells (Life Technologies Ltd.) by heat shock. Cells were grown on LB agar plates supplemented with spectinomycin (100 µg/ml). Colonies were selected and cultured, and extracted plasmids were sequenced with primers P03 and P04 to confirm the correct assembly of pOPARA2.

### Cloning into pOPARA1, pOPARA2 and pOPIN-GG

The coding sequence of *E. coli* maltose binding protein (MBP, (Duplay et al., 1984)) was PCR-amplified with primers that introduced flanking overhangs (5’ AATG and 3’ TTCG) and *BsaI* restriction sites into the PCR product (see Table 1 for details of the primers). The amplicons were subsequently combined with a level 0 C-terminal 6xHis tag module pICSL50025 (Addgene plasmid # 174589) and either pOPARA1, pOPARA2 or pPGC-C (Addgene plasmid # 174580, (Bentham et al., 2021)) for Golden Gate assembly in a digestion-ligation reaction as described in (Engler et al., 2008). Reaction products were used to transform chemically competent *E. coli* DH5α cells (Life Technologies Ltd) by heat shock. Positive (white) colonies were selected and cultured, and extracted plasmids were sequenced using primers P01 and P02 (Table 1) to confirm the correct assembly of the constructs.

### Production and purification of test proteins from pOPARA1, pOPARA2 and pOPIN-GG

Expression constructs were used to transform chemically competent *E. coli* BL21(DE3) cells by heat shock. Single colonies were used to inoculate 10 ml L media supplemented with appropriate antibiotics (carbenicillin or spectinomycin; final concentration of 100 µg/ml). Following overnight incubation at 37 °C with agitation, cultured cells were added to 2 L Erlenmeyer flasks containing 500 ml L media supplemented with appropriate antibiotics to a starting OD_600_ of 0.01-0.02. Cultures were incubated at 37°C with agitation until an OD_600_ of 0.5-0.7 was reached, then expression was induced with L-arabinose (pOPARA1 or pOPARA2 transformants, final concentration of 0.2%) or IPTG (pPGC-C, final concentration of 1mM). Cultures were incubated for a further ∼16 hours with agitation at 18 °C. Cells were harvested by centrifugation and resuspended in lysis buffer (50mM Tris pH 8.0 at 4°C, 500mM NaCl, 5% glycerol, 50mM glycine, 30mM imidazole, cOmplete EDTA-free protease inhibitor cocktail) at a ratio of 5ml lysis buffer per gram wet weight of pellet. Resuspended cells were lysed by sonication (Vibra-Cell™ sonicator with a 13 mm probe) and the cell lysate was clarified by centrifugation (16,500 x g for 30 minutes at 4 °C). The supernatant (soluble fraction) was applied to a 1ml HisTrap FF column (Cytiva) pre-equilibrated in binding/wash buffer (50mM Tris pH 8.0 at 4°C, 500mM NaCl, 5% glycerol, 50mM glycine, 30mM imidazole). Following sample application, the column was washed with 5CV of binding/wash buffer, and the bound protein eluted with 3CV elution buffer (50mM Tris pH 8.0, 500mM NaCl, 5% glycerol, 50mM glycine, 500mM imidazole). Samples were visualised by SDS-PAGE using 12% TruPAGE precast gels and the associated 4X LDS sample buffer and 1X TEA-Tricine running buffer (Sigma-Aldrich). 5 µl Spectra™ Multi-Color Broad Range Protein Ladder (Thermo-Scientific) was loaded alongside the samples. Gels were stained with Instant Blue™ (Expedeon).

## Acknowledgements

We thank H. Peter van Esse for critical reading of the manuscript. We thank Sylvestre Marillonnet for making pAGM1299 (Addgene plasmid # 47988 ; http://n2t.net/addgene:47988 ; RRID:Addgene_47988)available on Addgene. We thank Mark Banfield for making pPGC-C (Addgene plasmid # 174580; http://n2t.net/addgene:174580; RRID:Addgene_174580) and pICSL50025 (Addgene plasmid # 174589; http://n2t.net/addgene:174589; RRID:Addgene_174589) available on Addgene.

This work was supported by 2Blades.

## Supplemental data files

Supplemental data 1: GenBank format file for pOPARA1 Supplemental data 2: GenBank format file for pOPARA2

## Data availability

The pOPARA vectors are available from Addgene (pOPARA1: Addgene ID #200416, pOPARA2: Addgene ID #200417).

## Notes

https://www.addgene.org/200416/

https://www.addgene.org/200417/

### Competing Interest Statement

The authors have declared no competing interest.

